# Temporal expectation triggers competition in working memory that leads to forgetting

**DOI:** 10.64898/2026.03.26.714304

**Authors:** Ziyi Duan, Zhang Ziyao, Jarrod A Lewis-Peacock

## Abstract

Working memory (WM) provides a flexible but capacity-limited workspace for maintaining information over short intervals, whereas long-term memory (LTM) serves as a vast and enduring repository for preserving information over extended periods. Decades of research suggest that they are two distinct yet connected systems that together enable adaptive behavior. The link between WM and LTM may not be straightforward, however, as recent evidence has shown that activation-dependent competition among items in WM can weaken their representations in LTM. In the current study, we examined how dynamic competition among items for limited WM resources affects their retention in LTM. We induced competition between items by manipulating temporal expectations in a WM task with either a short (1 s) or a long (4 s) memory delay. Human participants (N = 20) initially prioritized items expected to be tested early, but shifted their priority to items expected to be tested later when the early test did not occur. Using electroencephalography (EEG) and multivariate pattern analysis (MVPA), we tracked the dynamic fluctuations in WM contents based on expected task relevance across the delay window. We linked these temporal profiles during WM with the long-term recognition performance of each item and found that forgetting was associated with a marked decrease in neural evidence for items deemed no longer relevant during the later delay period. These results demonstrate that WM representations fluctuate with temporal expectations and that the de-prioritization of items during WM maintenance is what drives their long-term forgetting.

## Introduction

Working memory (WM) is a capacity-limited online system that maintains and manipulates information over brief periods to support goal-directed behavior (A. Baddeley, 2003; A. D. Baddeley, 1986). In contrast, long-term memory (LTM) stores vast amounts of information over extended periods (Brady et al., 2008; Cowan, 2008; Ebbinghaus, 1913). Historically, these systems were viewed as neurally distinct: WM was linked to persistent prefrontal cortex (PFC) activity (Curtis & D’Esposito, 2003; Fuster & Alexander, 1971), while LTM was associated with the medial temporal lobe and hippocampus (Squire & Zola-Morgan, 1991).

However, emerging evidence highlights shared mechanisms between WM and LTM. Behavioral studies have shown that longer WM maintenance improved later LTM recognition (Hartshorne & Makovski, 2019; Souza & Oberauer, 2017). Moreover, emerging neural evidence suggests that the hippocampus also plays a crucial role in WM storage (Borders et al., 2022; Daume et al., 2024; Goodrich et al., 2019; Jeneson & Squire, 2012; Kamiński et al., 2017; Yonelinas, 2013).

Complementing this view, sensory cortical regions recruited during WM maintenance also appear to carry feature-specific mnemonic representations that evolve with task demands (Kiyonaga & Serences, 2025), suggesting that the neural substrate of WM storage extends beyond the hippocampus to include the very sensory areas that initially encoded the information. These findings suggest that neural activations during WM formation and maintenance could affect long-term traces in LTM.

One possibility is that the sustained maintenance of information in WM gradually strengthens LTM traces, thus predicting a linear relationship between WM and LTM (Hebb, 2012). Many human neuroimaging and neurophysiology studies provided evidence supporting this relationship. For example, previous functional magnetic resonance imaging (fMRI) studies found that the magnitude of neural activity in related brain regions during WM predicted subsequent LTM performance (Davachi et al., 2001; Ranganath et al., 2005; Schon et al., 2004). However, these studies often involved single items for a fixed delay interval, leaving it unclear how competition for limited WM resources affects long-term retention.

WM is known for having limited capacity (Awh et al., 2007; Luck & Vogel, 2013; Vogel et al., 2001). If persistent activity in WM is essential for long-term storage, competition amongst multiple items for limited WM resources could play a crucial role in determining which items are remembered or forgotten. Unlike prior models that posit a linear relationship, the non-monotonic plasticity hypothesis (NMPH) posits a U-shaped relationship between memory activation and learning: little to no activation does not alter memory strength; strong activation increases memory strength; and, critically, moderate levels of activation weaken memory (Detre et al., 2013; Newman & Norman, 2010; Ritvo et al., 2019). The NMPH is modeled on principles of memory retrieval and also predicts how competition among items in WM affects their long-term retention (Lewis-Peacock & Norman, 2014). Specifically, when two items compete for WM resources, for example, during a transition of task priorities, items with the highest level of activation should be strengthened, items with weak activation should not be impacted, and items with a moderate level of activation should be weakened, leading to forgetting.

In the current study, we tested how temporal expectations about an item’s task relevance affect its neural representation in WM (van Ede et al., 2017) and its retention in long-term memory. We induced competition between two memory items by associating each with a short (1 s) or long (4 s) probe interval, signaling whether it was expected to be probed early or late, but only one item was probed on each trial. This created a dynamic task environment in which prioritization amongst two items in working memory can switch depending on how much time has elapsed during the memory delay. We hypothesized that participants would initially prioritize the item expected to be tested early and, if the early test did not occur, would switch to prioritizing the other item expected to be tested late. This shift in prioritization over time could induce different levels of competition among memory items during the WM delay, leading to long-term forgetting.

## Methods

### Participants

Twenty-six healthy human volunteers participated in the study. The sample size was determined based on prior related studies (Detre et al., 2013; Kim et al., 2014; Lewis-Peacock & Norman, 2014). All participants had normal or corrected-to-normal vision. All participants provided written informed consent before participation. The experimental protocols were approved by the University Committee on Activities involving Human Subjects at the University of Texas at Austin. Data for two participants were not collected due to technical issues, and data from four participants were excluded due to excessive noise, leaving 20 participants for the final analyses (11 males, 9 females; mean age = 23.1, SD = 7.1).

### Stimuli

A large collection of face (Tottenham et al., 2009), scene (Konkle et al., 2010), and object (Brady et al., 2008) stimuli was gathered through various online and in-house sources. A subset of these stimuli was chosen for this experiment based on memorability ratings from a stimulus evaluation experiment conducted through Amazon.com’s Mechanical Turk (Lewis-Peacock & Norman, 2014). The final stimulus set consisted of 86 grayscale images of each category for the Phase 1 localizer task (36 for memory targets, 50 for memory lures). For the Phase 2 WM task, the final stimulus set consisted of 194 grayscale images of faces and 194 grayscale images of scenes (144 memory targets and 60 memory lures). Another 8 images of faces and scenes were added for practice at the beginning of each block. The final Phase surprise LTM task consisted of a new set of 24 grayscale face images and 24 grayscale scene images as memory lures. Unique subsets of stimuli were used in each of the three phases of the experiment, and no target stimulus was ever reused as a target in another trial within a phase. The eight-picture probe display on each Phase 1 and Phase 2 trial was constructed by randomly sampling (with replacement across trials) from the set of lures. The assignment of stimuli to experimental phases and (within phases) to the target and lure conditions was done randomly for each participant.

### Task and Procedure

Each participant performed a three-phase task (see **Fig. 1**). Phase 1 served as a localizer task to train a categorical classifier of patterns of EEG voltage amplitudes. Phase 2 consisted of a WM task that created competition between two items based on temporal expectations of their task relevance. We acquired EEG data while participants performed the Phase 1 and Phase 2 tasks. Lastly, Phase 3 was a surprise LTM task that only included behavioral tests.

**Fig. 1.**
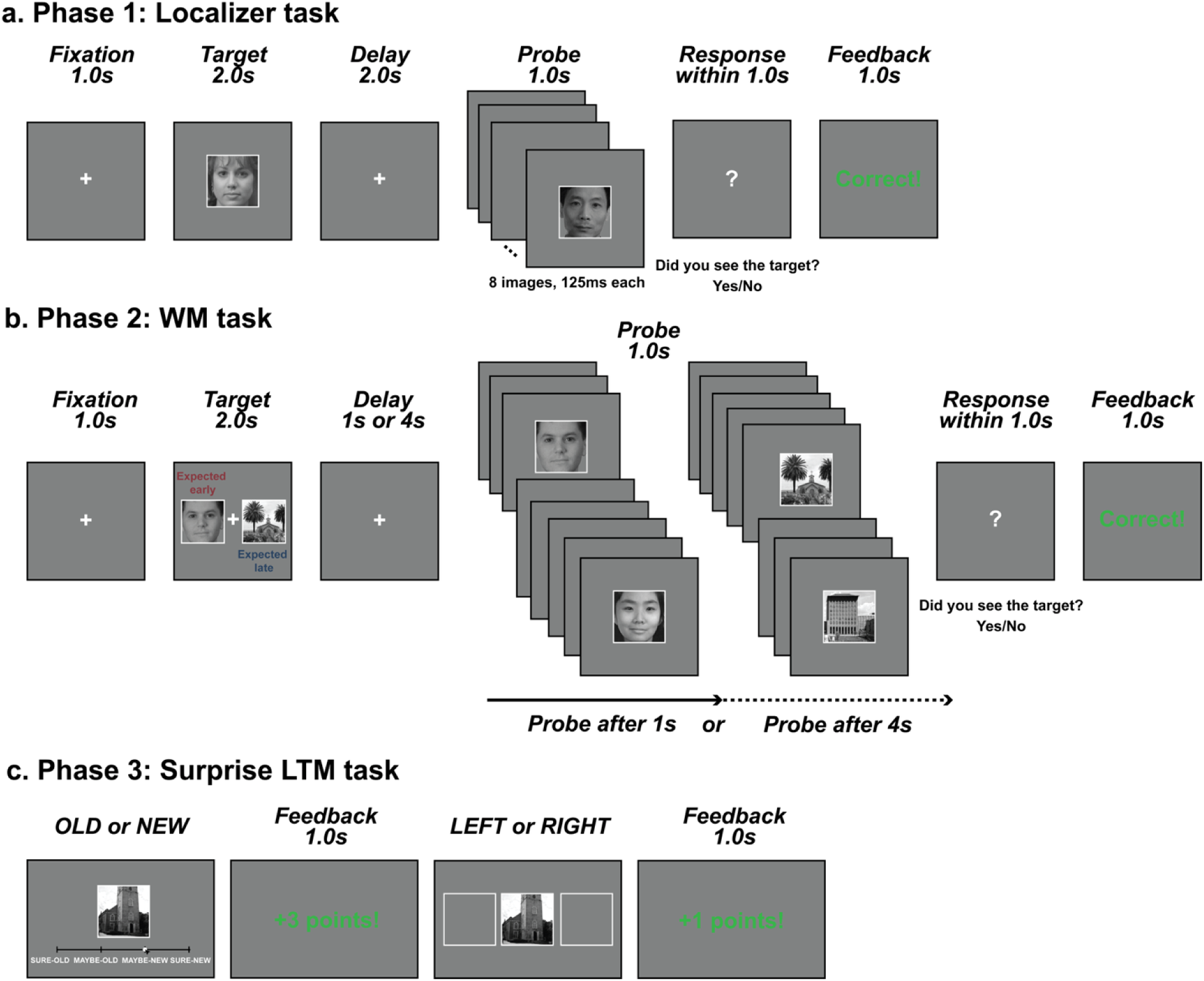
Task procedures. a). The localizer task was used to train a categorical classifier. Participants needed to detect a target item in an RSVP of eight images from the same category (face, scene, or object). b). In the WM task, two target images (face and scene) were presented. After a short (1 s) or a long (4 s) delay, one image was tested. If probed after a short delay, the face was tested; whereas if probed after a long delay, the scene was tested. c). Participants completed a surprise LTM task at the end of the experiment to test the influence of WM competition on long-term retention. Participants first rated their confidence in recognizing an image, then recalled the image’s location (left/right). Face and scene images were chosen from open datasets (Konkle et al., 2010; Tottenham et al., 2009) used in this study.

### Phase 1: Localizer task

We used a perceptual localizer task to train a categorical classifier to distinguish faces, scenes, and objects. In total, we had three blocks, each for a different category. Faces and scenes were used in the next two phases, and objects served as a baseline category to help the decoder decorrelate the face and scene categories. Each block consisted of 36 trials. Each trial began with a 1s central fixation, followed by the presentation of the target for 2s, and then a 2s delay.

Participants were instructed to remember the target item and complete a detection task after the delay. All probe displays consisted of a rapid serial visual presentation (RSVP) of eight images (from the same category as the target) presented in the center of the screen (125ms each). The target reappeared in the probe stream on half of the trials (in serial positions 3, 4, 5, or 6, chosen randomly), and lures were selected at random from a set of 50 same-category images that were never presented as targets. Participants were asked to report within 1 s whether the target was in the series. After their response, feedback was presented for 1s (correct, incorrect, or late).

### Phase 2: Working memory task

During encoding, two images (one face, one scene) were presented for 2 s on the left and right sides of the fixation. The location of the face and the scene were randomly assigned for each trial and counterbalanced within each block. Participants were required to remember both images, but only one was probed after one of two possible delays (1 s or 4 s). The key manipulation was that, when probed after a short delay (1 s), the face was tested; when probed after a long delay (4 s), the scene was tested. In other words, faces were expected to be tested early, while scenes were expected to be tested late. By linking the face to a short delay and the scene to a long delay, we expected participants to prioritize face representation at the beginning of trials and, if the early test never appeared, to reprioritize and shift to scene representation. During the probe period, participants had to determine if the probed image was present in an RSVP of eight images from the same category, structured as in the localizer task.

In total, there were three blocks, each with 48 trials. In blocks 1 and 3, we set the probabilities of short- and long-delay trials to 25% and 75%, respectively. The asymmetry between short- and long-delay trials was intended to increase statistical power to test our critical hypothesis following the shift in prioritization in the long-delay trials. In block 2, we set the probability to 50% for both conditions to test for possible effects arising from the unbalanced design.

Participants were informed about the category expectations (face: short; scene: long) and the short/long testing probability (25/75 or 50/50) at the beginning of each block and completed 8 practice trials before the real task.

### Phase 3: Surprise long-term memory task

At the end of the experiment, participants were asked to complete a surprise recognition memory test. This test was restricted to items that were exposed but not tested during the WM task. That is, for short-delay trials, only expected-late items (scenes) were tested, whereas for long-delay trials, only expected-early items (faces) were tested. In total, we had 144 old items (96 faces and 48 scenes) together with 24 new faces and 24 new scenes as lures. For each trial, participants were asked to rate their confidence that an item was old/new in an image recognition task. The confidence scale had 4 levels (1 = sure old, 2 = maybe old, 3 = maybe new, 4 = sure new). For old trials, participants were also asked to determine the target’s location on the screen (left or right) during the WM task. For each image, we provided feedback on their performance for 1s, which was tied to a point-getting system for motivation. Specifically, if participants’ answers to the old/new task were correct, they earned 3 points for a "sure" answer and 1 point for a "maybe" answer. On the contrary, if their answers were wrong, they lost 3 points for a sure answer and 1 point for a maybe answer. For the location task, participants received an additional point for correct responses and no points for incorrect responses. Every 100 points was converted into $0.50 of bonus pay.

### EEG Acquisition

The EEG signal was acquired using a 64-channel Biosemi ActiveTwo system, with electrodes arranged according to the International 10–20 system. Data was recorded at 1,024 Hz. Additional electrodes were placed on the mastoids, below the right eye, to the left of the left eye, and to the right of the right eye.

### EEG Preprocessing

EEG data were analyzed using Python and the MNE software (Gramfort et al., 2014). During data preprocessing, we first referenced raw data to the average of the left and right mastoid measurements. Then, we removed line noise using the notch filtering provided by MNE. The signal was then band-pass filtered between 0.01 and 80Hz. If too many slow drifts were detected, 0.2Hz was used as a low cutoff. Next, we removed corrupted data spans by detecting eye blinks using the EOG channels. During the data epoch, we first rejected epochs by both setting the maximum and minimum peak-to-peak amplitude to be 100 and 10^-5^µV. Then, after the data epoch, we conducted visual inspections of the signal’s variance across trials and channels.

Finally, we applied baseline correction using signals 200ms before the start of each trial and downsampled the data to 500Hz (for similar preprocessing approaches, see (Bae & Luck, 2018)).

### EEG Data Analysis

#### Training image category decoders

Image category (object, face, and scene) classifiers were trained using data from the localizer task. We focused on the later stage of encoding (1-2 s post-stimulus onset) to capture reliable sensory representations for each image category. We were unable to detect reliable representations during the localizer task’s delay window; therefore, we used only the encoding window for training. We used a support vector machine (SVM) to classify the target item’s category based on the spatial distribution of EEG voltages across electrodes. This model was implemented using the SVC function from the sklearn package in Python (Pedregosa et al., 2012). EEG signals were averaged over the 400-600 ms window after image onset, when visually evoked responses were most reliable. To optimize the classifier for each participant, a grid search was performed on the localizer data to select the best C regularization parameter (range: 0.0001–10000).

#### Capturing working memory fluctuations

We applied the trained category classifiers to track image representations during the main WM task. Each non-overlapping time point (10 ms) in the main task phase was treated as a test sample. For each participant, we used the same set of electrodes identified during the training phase. The classifier output prediction probabilities for each of the three image categories, which we interpreted as evidence for each category. For analysis, we focused only on the relevant face and scene categories. Evidence for object categories was consistently at or below the empirical chance level of 0.33, as expected, and was therefore excluded from the analyses.

We used linear regression models to examine changes in category evidence over time. The outcome variable was category evidence for the face and scene categories, and the sole predictor was time. Positive beta values indicated an increasing trend in category evidence over time, while negative beta values indicated a decreasing trend. This analysis was conducted separately for early (0-1 s after the delay onset) and late (1-4 s after the delay onset) time windows, as our manipulation of testing probabilities was expected to shift temporal expectations.

#### Linking working memory fluctuations to long-term memory performance

Here, we focused on linking neural evidence for the untested face image during the long-delay WM trials with its LTM performance. First, we separated face images into remembered and forgotten items based on the LTM recognition task. Given insufficient trials across all four confidence levels, we collapsed trials with “sure” and “unsure” responses and defined remembered items as those with the response “old” and forgotten items as those with the response “new”. Then, we calculated the category evidence for the face and scene for both the remembered and forgotten trials in the WM task. Moreover, we calculated the category evidence difference between scene and face for each trial, with positive values indicating stronger neural evidence for scene images than for face images. Then, we compared the category evidence difference between remembered and forgotten trials. Finally, we averaged the category evidence difference during the early (0-1 s after delay onset) and late (1-4 s after delay onset) windows for statistical testing.

### Statistics

We used repeated-measures ANOVAs to compare decoding performance across image categories (face and scene) and time windows (early and late). Planned paired-sample t-tests were conducted to assess differences between conditions, and one-sample t-tests were used to evaluate decoding performance against chance levels. All analyses were conducted in Python using the pingouin statistical package. Bayes factors in favor of the alternative hypothesis (BF10) were reported, with values between 1 and 3 indicating anecdotal evidence, and values between 3 and 10 reflecting moderate evidence. Sidak correction was applied to adjust for multiple comparisons (Sidak, 1967), and corrected p-values were reported. All t-tests were two-sided unless otherwise noted in the main text.

Because the testing probability for Block 2 (50/50) differs from that for Blocks 1 and 3 (25/75), we reported EEG results excluding Block 2 in the main text. Results, including Block 2, are included in the supplementary material (**Sfig. 2, 3b, 4b, and 6**), showing a similar pattern but weaker statistics.

**Fig 2.**
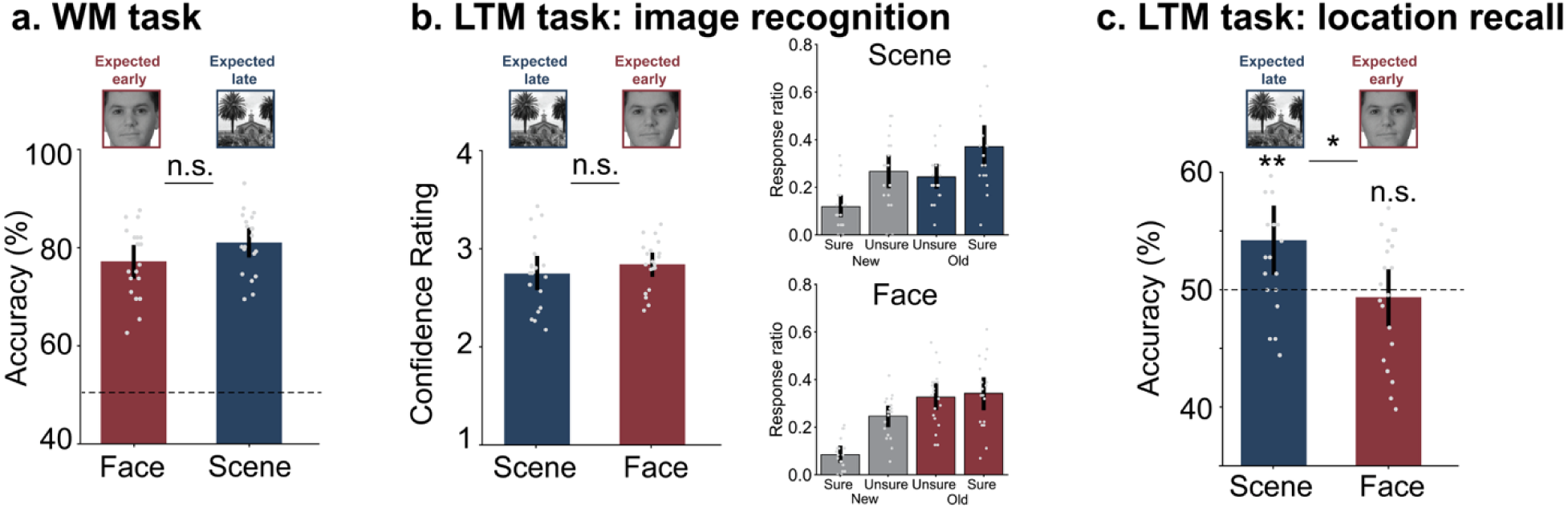
Behavioral performance for WM and LTM tasks. a). Averaged response accuracy for the RSVP WM task. Face images were tested in short-delay trials, while scene images were tested in long-delay trials. b). Averaged confidence rating (left) and response ratio for each confidence level (right) in the surprise LTM image recognition task. For short-delay trials, scene images that were not tested in the WM task were tested in the LTM task. For long-delay trials, face images that were not tested in the WM task were tested in the LTM task. c). Averaged response accuracy for location recall for face and scene images tested in the LTM task. Error bars represent 95% confidence intervals. * indicates corrected p < .05; ** indicates corrected p < .01; n.s. is not significant. Face and scene images were chosen from open datasets (Konkle et al., 2010; Tottenham et al., 2009) used in this study.

**Fig 3.**
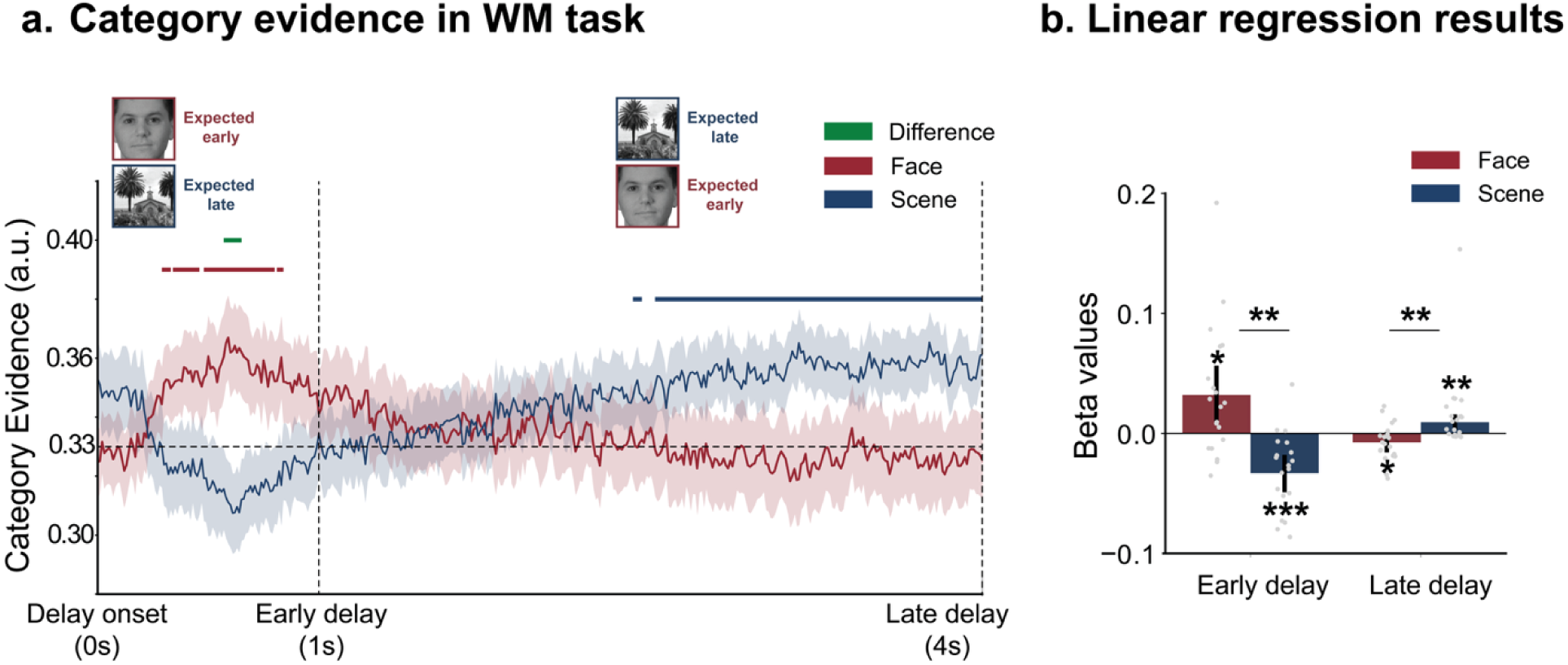
Temporal Fluctuations of Working Memory Representations. a). Temporal dynamics of category evidence for face and scene images in the WM task for the long-delay trials. Red horizontal lines indicate time points where face evidence was significantly above the empirical chance level of 0.33; blue lines indicate significant scene evidence; green lines indicate time points where face and scene evidence differed significantly. The shaded area represents standard errors. b). Beta values from linear regression models predicting category evidence from time, separately for the early (0–1 s) and late (1–4 s) delay windows. Error bars represent 95% confidence intervals. * indicates corrected *p* < .05; ** indicates corrected *p* < .01; *** indicates corrected *p* < .001. Face and scene images were chosen from open datasets (Konkle et al., 2010; Tottenham et al., 2009) used in this study.

**Fig 4.**
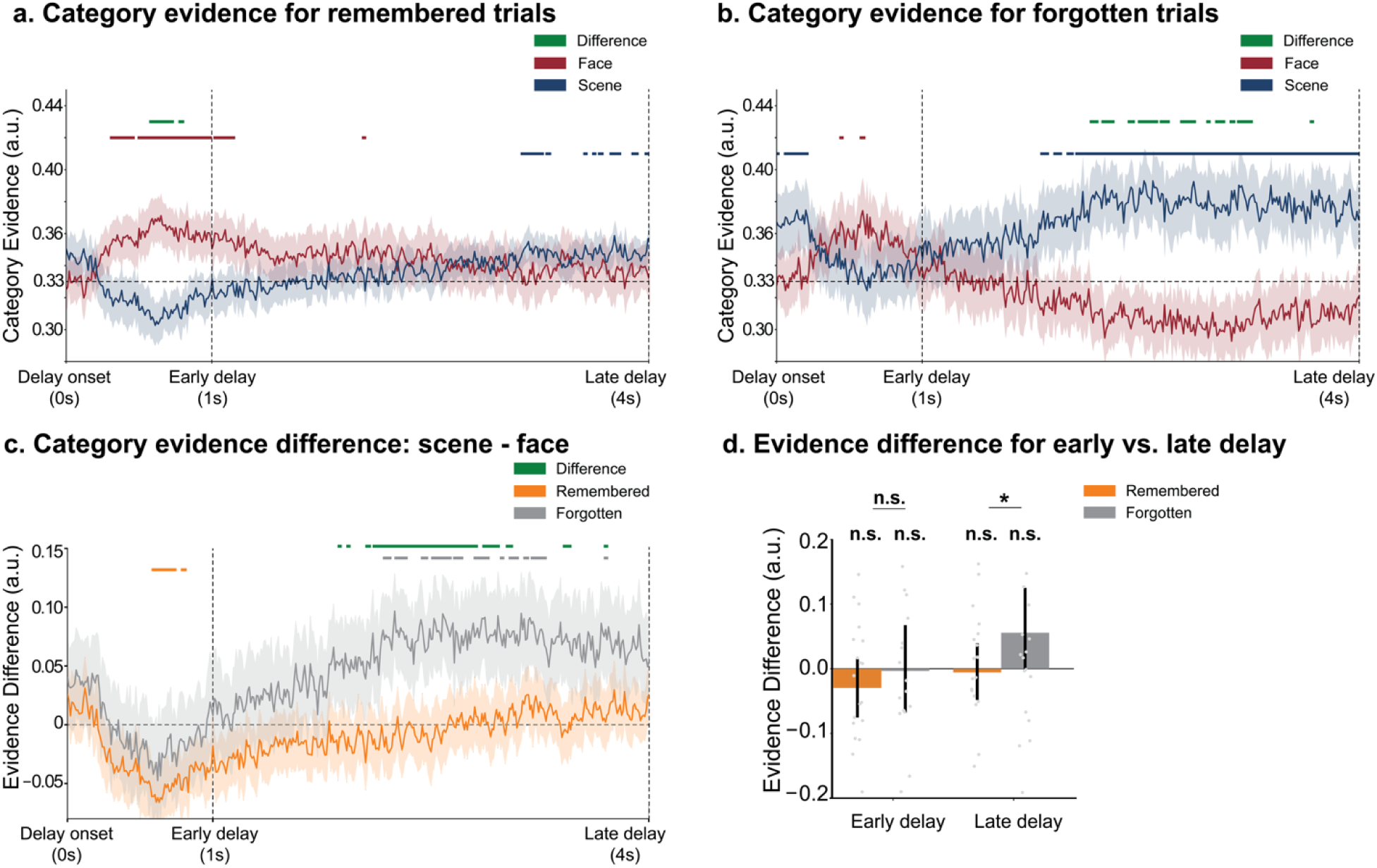
Differentiated temporal profiles for remembered and forgotten trials. a). Temporal dynamics of face and scene category evidence for trials in which face images were later remembered in LTM. b). Temporal category evidence for trials in which face images were later forgotten. Red horizontal lines indicate time points where face evidence was significantly above the empirical chance level (0.33); blue lines indicate significant scene evidence; green lines indicate significant differences between face and scene evidence. c). Differences in category evidence between face and scene for remembered and forgotten trials. Forgotten trials showed a larger difference between face and scene category evidence than remembered trials during the late delay window. Orange horizontal lines indicate time points where category evidence difference between face and scene was significantly above 0 for the remembered trials; grey lines indicate significant evidence difference for forgotten trials; green lines indicate significant differences between remembered and forgotten trials. The shaded area represents standard errors. d). Averaged category evidence difference for early (0–1 s) and late (1–4 s) delay windows. Error bars represent 95% confidence intervals. * indicates corrected *p* < .05; n.s. is not significant.

## Results

### Behavioral performance

Here, we compared WM and LTM task performance based on whether the probed item was a face or a scene in each task. For example, when the face image (expected early) was probed after a short delay during the WM task, only the scene image (expected late) from the same WM trial would be tested in the LTM task. In the WM task, there was no significant difference in accuracy when detecting the face image after a short delay compared to detecting the scene image after a long delay (see **Fig. 2a**), *t*(19) = 2.03, *p* = .056, *d* = 0.58, *BF10* = 1.26. To examine the LTM fate of competitive items in WM, we conducted two types of memory tests: 1) An image recognition test, where participants rated their confidence of recognizing a test image as old/new. 2) An incidental source memory test, where participants recalled the initial presentation location of an old image. First, we calculated the averaged confidence rating for the image recognition task and found no difference between the face and scene images, *t*(19) = 1.57, *p* = .133, *BF10* = 0.66 (**Fig. 2b)**. For the location recall task, we found better location recall accuracy for the scene images compared to the face images (**Fig. 2c**), *t*(19) = 2.55, *p* = .020, *d* = 0.83, *BF10* = 2.94. Participants performed above chance level for scene images [*t*(19) = 2.98, *p* = .008, *d* = 0.67, *BF10* = 6.43], but not face images [*t*(19) = -0.54, *p* = .593, *BF10* = 0.27]. We discuss the potential reason for better location recall for the scene in the Discussion. To link LTM outcomes to neural activation of the representations in WM, we correlated behavioral performance in the image recognition task with trial-by-trial category-based pattern classification results from the delay-period of the WM task. But first, we will describe how these pattern classifiers performed.

### Training image category decoders

Three-class category decoders were trained using One-vs-Rest (OVR) logistic regression, where we trained three separate binary logistic regression classifiers (one per class), each distinguishing "class k" from "all other classes." At test time, the three classifiers return a probability for selecting their trained class. This was done separately for each participant using EEG data from the encoding period (1-2 s) of the localizer task. A repeated-measures ANOVA (predicted category by actual category) was conducted to assess decoder reliability, and showed a significant interaction (**Sfig. 1**), *F*(4,76) = 22.06, *p* < .001, 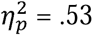. The prediction probabilities were significantly higher for the correct target category than for the other categories across objects, faces, and scenes (objects: *F*(2,38) = 7.02, *p* = .002, 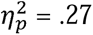; faces: *F*(2,38) = 12.61, *p* < .001, 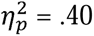; scenes: *F*(2,38) = 20.30, *p* < .001, 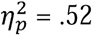), indicating that the trained decoders reliably distinguished EEG patterns associated with processing each image category.

### Temporal expectations modulate fluctuations in working memory representations

We used trained category decoders within each individual to track dynamic changes in WM representations during the memory delay period. Temporal expectations were manipulated so that face images should be prioritized early during the delay period (0-1 s into the delay), whereas scene images should be prioritized later (1-4 s into the delay). Here, we examined fluctuations in neural classification evidence during long-delay trials, in which participants needed to reprioritize from face to scene images over time (results for short-delay trials are in supplementary, see **Sfig. 4**). Decoding results revealed that face representations increased early in the delay and then declined, while scene representations ramped up later in the delay period (**Fig. 3a**, for the full timecourse, see **Sfig. 3**).

To quantify these dynamics, we fitted simple linear models to assess whether face and scene representations increased or decreased during the early and late delay windows (**Fig. 3b**). A significant interaction was found between time window and stimulus category, *F*(1,19) = 15.08, *p* = .001, 1]^2^ = .29. During the early delay window, neural evidence for face images showed a reliably positive slope, *t*(19) = 2.64, *p* = .016, *d* = 0.59, *BF10* = 3.44, while neural evidence for scene images significantly decreased, *t*(19) = -4.52, *p* < .001, *d* = 1.01, *BF10* = 130.47. The slope was significantly stronger for face images than for scene images, *t*(19) = 3.70, *p* = .002, *d* = 1.45, *BF10* = 25.47. These results suggested that participants initially prioritized the face image during the early delay window. Interestingly, these patterns reversed in the late delay window.

Category evidence values for scene images increased reliably, *t*(19) = 3.78, *p* = .001, *d* = 0.84, *BF10* = 29.54, while for face images they decreased, *t*(19) = -2.11, *p* = .048, *d* = 0.47, *BF10* = 1.44. This change in category evidence across the delay was greater for scene images than for face images, *t*(19) = 3.12, *p* = .006, *d* = 1.23, *BF10* = 8.28. This transition indicated a switch of prioritization from the face to the scene image during the late delay window.

### Subsequent forgetting is linked to switching between working memory representations

To examine how LTM outcomes relate to representational fluctuations in WM, we focused on long-delay trials in which participants initially prioritized the face image but later shifted prioritization to the scene image. We hypothesized that this switch in representational priority, from faces to scenes, might lead to competition between these items, contributing to the forgetting of the de-prioritized face items. To test this, we separated trials by whether the face images were later remembered or forgotten (see Methods). First, we visualized the category evidence for both face and scene images in the WM task, separately for the remembered (**Fig. 4a**) and forgotten (**Fig. 4b**) trials. In both cases, we observed that the category evidence for face peaked during the early delay and declined thereafter, while the category evidence for the scene ramped up in the late delay (for the full timecourse, see **Sfig. 5**). Importantly, we noticed a much larger difference in the category evidence values for the face and scene in the forgotten trials than in the remembered trials. To directly compare the difference, we calculated the trial-wise category evidence difference between face and scene, and compared it between trials where the face was remembered vs. forgotten (**Fig. 4c**). We observed a significantly higher category evidence difference for remembered than forgotten trials during the late delay time window.

During the early delay window, no reliable differences existed between the category evidence values for remembered vs. forgotten trials [remembered: *t*(19) = -1.31, *p* = .206, *BF10* = 0.49; forgotten: *t*(19) = -0.10, *p* = .921, *BF10* = 0.23; remembered-forgotten: *t*(19) = -0.97, *p* = .342, *BF10* = 0.35]. However, during the late delay window, forgotten trials showed stronger category evidence differences than remembered trials [remembered: *t*(19) = -0.26, *p* = .801, *BF10* = 0.24; forgotten: *t*(19) = 1.70, *p* = .106, *BF10* = 0.78; remembered-forgotten: *t*(19) = -2.15, *p* = .045, *d* = 0.89, *BF10* = 1.52]. These results suggest that a more complete representational switch from the face to the scene was associated with greater forgetting of the de-prioritized item (i.e., face).

Finally, we tested whether forgetting faces in the LTM task was associated with higher accuracy on the competitor items (i.e., scenes) tested in the WM task. This could reflect a competitive dynamic between the items in WM that trades off better memory for the task-relevant item in the short term for worse memory for the task-irrelevant item in the long term. However, this analysis revealed no difference in WM accuracy for the scene on trials in which the face was later remembered vs. forgotten [hit rate: *t*(19) = 0.66, *p* = .517, *BF10* = 0.28].

## Discussion

The activation of representations in WM fluctuates over time due to multiple factors, including attentional priority, interference among co-activated items, and – as demonstrated here – expectations of their task relevance. In the current experiment, we manipulated temporal expectations about items held in WM to create dynamic competition between them and tested the long-term consequences. Using MVPA applied to EEG data, we tracked real-time fluctuations in the prioritization of WM contents as participants flexibly redirected their internal attention based on their expectation about when each item would be tested. We found that neural representations produced by the dynamic switch in WM prioritization predicted the fate of de-prioritized items in LTM. Specifically, long-term forgetting of the de-prioritized face images was associated with a marked decrease in neural evidence for the face images *and* a concurrent increase in neural evidence for scenes – precisely when faces were no longer relevant to the WM task. Together, these findings demonstrate that the neural fate of an item during WM maintenance — shaped here by temporal expectation — can determine whether it is ultimately remembered or forgotten, linking the dynamics of WM prioritization directly to LTM outcomes.

### Multivariate EEG patterns track dynamic prioritization in WM

Many neuroimaging studies have revealed the dynamic and flexible nature of WM by manipulating task relevance across multiple items (Griffin & Nobre, 2003; Li et al., 2025; Myers et al., 2017). For example, in tasks that require a switch in prioritization, neural signals for WM items can be inactive and can be reactivated by retrospective cues, visual stimulation, or brain stimulation (LaRocque et al., 2013; Lewis-Peacock et al., 2012, 2015; Rose et al., 2016; Wolff et al., 2017). Moreover, recent studies have demonstrated the role of temporal expectation in establishing prioritization amongst items in WM in more dynamic settings (Jin et al., 2020; van Ede et al., 2017; Zokaei et al., 2019). They found that the switch in prioritization was associated with attenuation of contralateral alpha (8-14 Hz) oscillations. Although alpha lateralization reflects shifts in spatial attention, it provides no insight into how much information about the item is maintained in WM.

In the present study, we utilized machine learning to decode category-specific patterns of EEG activity for WM items associated with different temporal expectations – whether an item was relevant early or later in the memory delay. Importantly, we observed a dynamic switch in the prioritization of WM items across the early/late temporal threshold. During the encoding period across all trials, there was greater category evidence for scenes than for faces (**Sfigs. 3a**), indicating a bias toward encoding scenes over faces. This is likely due to our experimental design, which assigned a higher testing probability to scene images (75% in blocks 1 and 3) and always tested scenes after a long (4 s) delay whenever they were tested — meaning participants may have needed to form a stronger initial representation of scenes to withstand a period of de-prioritization before being reactivated and re-prioritized later in the delay. However, we observed greater category evidence for the face than the scene image during the early delay window, suggesting that participants prioritized the face image at the most relevant time period. Importantly, as time elapsed, neural evidence for the scene image revived – demonstrating that participants did not simply abandon the scene, but actively reinstated it as the likelihood of a delayed test grew.

We examined alpha lateralization in our WM task, but didn’t find any robust effect during the WM delay (see **Sfig. 7**). This is likely because the probed items during the response period were presented centrally, so the spatial location was not task-relevant. Overall, our results extend prior work by demonstrating how temporal expectations of task relevance shape the neural fate of competing items in WM over time.

### Different profiles of WM prioritization predict long-term forgetting

The main motivation of the current study was to test how dynamic competition among multiple items in WM influences their fate in LTM. We didn’t observe a difference in item recognition performance for scenes (when the face had been tested after a short delay) and faces (when the scene had been tested after a long delay), but we observed better location recall (left vs. right side of the screen) for scenes than for faces. There could be two reasons for enhanced incidental recall of scene locations. First, the number of scene images tested in the LTM task was much smaller than that of face images (48 scenes vs. 96 faces) because of unequal testing probabilities for short- and long-delay trials in the WM task. Therefore, reduced inter-trial interference may have benefited location recall for scene images. Second, scene images tested in the LTM task were drawn from short-delay WM trials, where face-scene competition was relatively brief – yet why reduced competition benefited location recall but not image recognition performance for scenes remains unclear and warrants future investigation.

To link LTM performance to neural competition during the WM task, we focused on long-delay trials in which participants initially prioritized the face but then shifted priority to the scene after no face test occurred – the most common trial type, comprising two-thirds of all trials. We separated these trials into “remembered” and “forgotten” bins based on LTM outcomes for the faces and examined category-specific EEG classifier evidence for faces versus scenes during the memory delay in the WM task. First, we found a positive, linear relationship between the WM activation level and subsequent LTM performance, such that remembered trials had stronger neural evidence for face images (as shown in **Fig. 4c**). These results supported the linear hypothesis and are consistent with previous findings that have linked stronger WM activation with better long-term retention (Daume et al., 2024; Davachi et al., 2001; LaRocque et al., 2015; Ranganath et al., 2005; Tozios & Fukuda, 2020). Second, forgotten face images were associated with stronger neural evidence for their competing scene images in WM, suggesting that de-prioritization involves an active reallocation of WM resources toward the newly relevant item rather than simply switching off the task-irrelevant one.

Our results bear a complex relationship to the NMPH (Detre et al., 2013; Newman & Norman, 2010; Ritvo et al., 2019), which predicts that moderate, competitive levels of activation for task-irrelevant items in WM should drive their long-term weakening and forgetting (Lewis-Peacock & Norman, 2014). At first glance, our data appear to invert this prediction: face images that were later *remembered* showed neural evidence comparable to that of the relevant scene image (**Fig. 4a**), suggesting sustained competition – conditions the NMPH associates with forgetting. Conversely, face images that were later *forgotten* showed markedly weaker neural evidence than the relevant item (**Fig. 4b**), suggesting little competition – conditions the NMPH associates with remembering. Rather than dismissing these findings as contradictory, however, we think they reveal important boundary conditions on when and how NMPH-like competition effects emerge.

Several factors may account for this discrepancy. First, competition in our task was driven by self-monitored temporal expectation rather than an explicit retro-cue, likely introducing greater trial-by-trial variability in the timing and completeness of prioritization shifts. Second, our effective competition window (∼ 3s post-switch) was substantially shorter than that of Lewis-Peacock & Norman (2014), who used an 8s window both before and after the priority switch – suggesting that NMPH-driven forgetting may require a longer accumulation of competitive activation. Third, EEG provides high temporal resolution for tracking dynamic WM fluctuations but is less sensitive to signals from medial and ventral temporal regions – including hippocampus – that are central to the LTM effects reported in prior fMRI work.

These discrepancies point to meaningful refinements for the NMPH. The hypothesis would benefit from explicitly incorporating temporal factors – particularly the duration and consistency of competitive activation – as modulators of the forgetting effect. More broadly, a persistent empirical challenge for NMPH-based predictions is defining what constitutes ‘moderate’ activation in any given experimental context, which makes it difficult to prospectively identify the conditions most likely to produce the nonmonotonic relationship between activation and memory strength. Our results underscore this challenge and suggest that future work should systematically vary the competition duration and measurement approach to better characterize the conditions under which WM competition translates into long-term forgetting.

## Conclusion

In sum, these findings advance our understanding of how the contents of WM are dynamically regulated and how those dynamics shape what is ultimately retained in LTM. By demonstrating that temporal expectations drive real-time competition between representations in WM — and that the neural consequences of that competition predict long-term remembering and forgetting — our results highlight a previously underappreciated mechanism by which WM actively shapes LTM. Rather than serving as a passive holding space, WM emerges here as a dynamic and selective gateway, where the ebb and flow of representational priority leaves a lasting imprint on memory. Temporal expectation, in particular, proves to be a potent and ecologically relevant driver of these dynamics, reflecting the kinds of prospective, goal-directed demands that characterize memory use in everyday life.

## Data availability statement

The processed EEG data and raw behavioral data generated in this study have been deposited in the Open Science Framework at https://doi.org/10.17605/OSF.IO/DGF8U. We also make all code used to analyze the EEG and behavioral data publicly available.

## Author contribution

J.A.L.-P and Z.D conceived of the idea and designed the experiment. Z.D and Z.Z collected the data. Z.D and Z.Z analyzed the data and visualized the results. Z.D and Z.Z. wrote the original draft of the manuscript. J.A.L.-P, Z.D., and Z.Z. reviewed and edited the final draft of the manuscript.

## Acknowledgments

We thank Dr. Elizabeth Lorenc for helping with the EEG equipment setup and data collection.

