## Supplementary for "Temporal expectation triggers competition in working memory that leads to forgetting"

**Supplementary Materials**

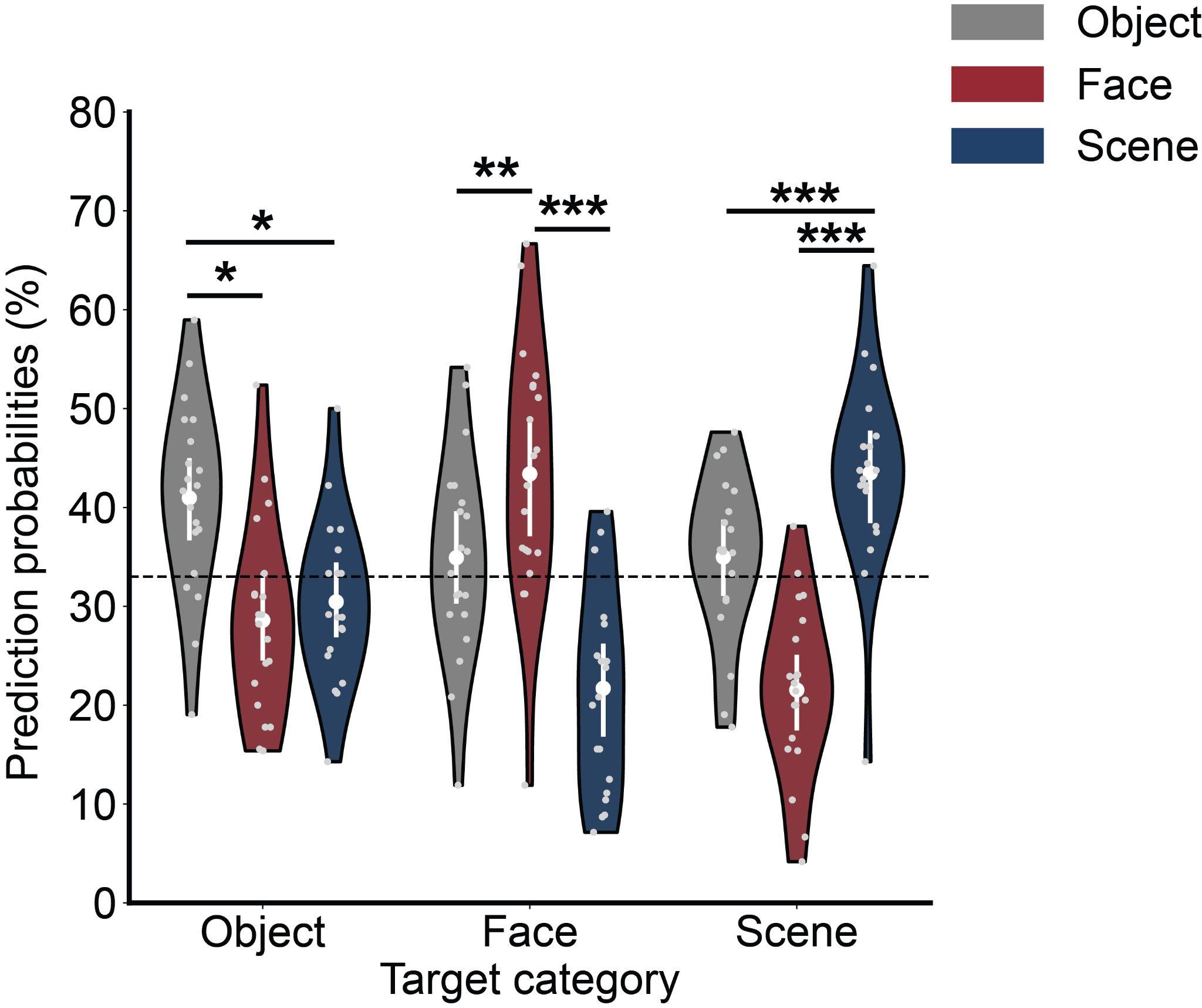

**Sfig 1.** **Decoding performance trained on the encoding window of the localizer task.** Prediction probabilities for the target categories were significantly higher than for non-target categories, demonstrating the reliability of the trained decoders. Error bars represent 95% confidence intervals. * indicates corrected *p* < .05; ** indicates corrected *p* < .01; *** indicates corrected *p* < .001.

To quantify the temporal dynamics of WM representations during the WM delay for all three blocks, we fitted simple linear models to assess whether face and scene representations increased or decreased during the early and late delay windows (**Sfig. 2**). A significant interaction was found between time window and stimulus category, *F*(1,19) = 14.70, *p* = .001, $\eta_{p}^{2}$ = .29. During the early delay window, neural evidence for face images showed a reliably positive slope, *t*(19) = 2.89, *p* = .009, *d* = 0.65, *BF10* = 5.41, while neural evidence for scene images significantly decreased, *t*(19) = -4.21, *p* < .001, *d* = 0.94, *BF10* = 70.40. The slope was significantly stronger for face images than for scene images, *t*(19) = 3.84, *p* = .001, *d* = 1.50, *BF10* = 33.46. During the late delay window, neural evidence for scene images increased reliably, *t*(19) = -2.18, *p* = .042, *d* = 0.49, *BF10* = 1.60, whereas for face images it decreased, *t*(19) = 3.16, *p* = .005, *d* = 0.71, *BF10* = 9.00. Besides, this increase was stronger for scene images than for face images, *t*(19) = -2.95, *p* = .008, *d* = 1.18, *BF10* = 6.00. These results are consistent with those reported in the main text, excluding Block 2.

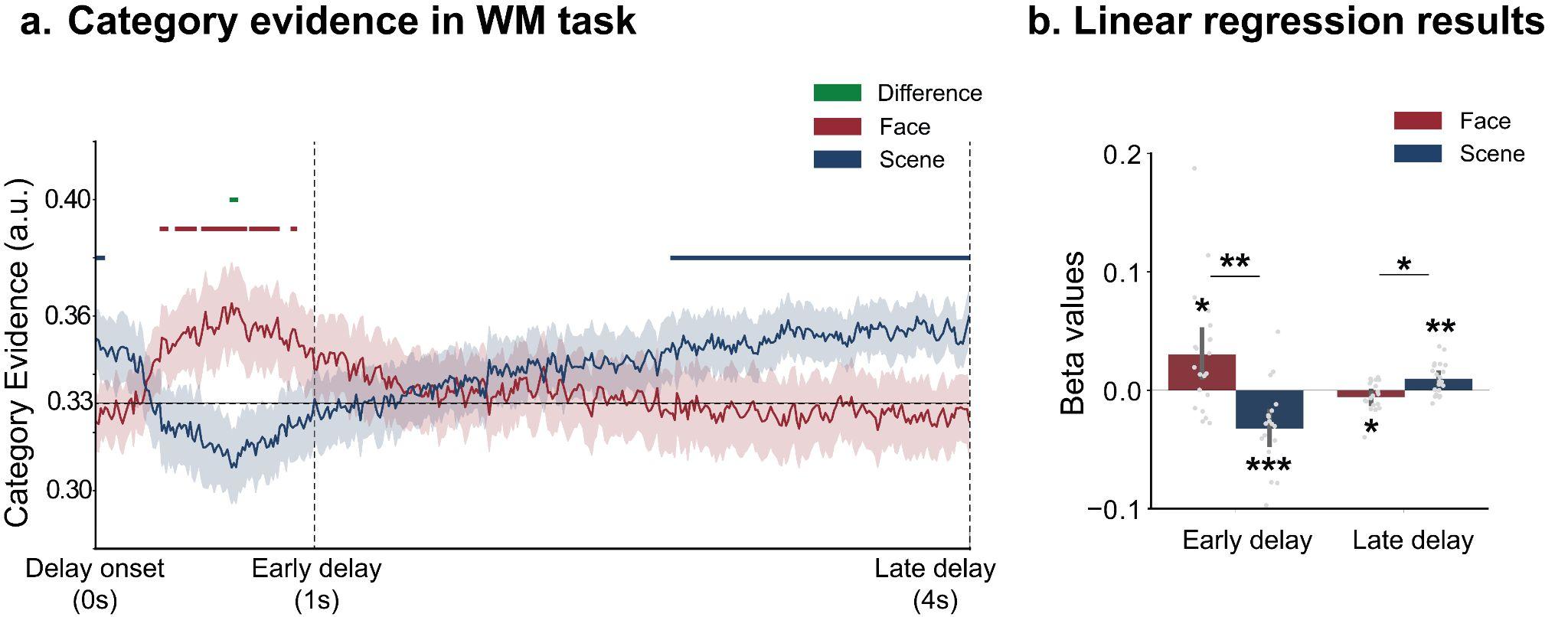

**Sfig 2.** **Temporal Fluctuations of Working Memory Representations for all three blocks.** a). Temporal dynamics of category evidence for face and scene images in the WM task for the long-delay trials. Red horizontal lines indicate time points where face evidence was significantly above the empirical chance level of 0.33; blue lines indicate significant scene evidence; green lines indicate time points where face and scene evidence differed significantly. The shaded area represents standard errors. b). Beta values from linear regression models predicting category evidence from time, separately for the early (0–1 s) and late (1–4 s) delay windows. Error bars represent 95% confidence intervals. * indicates corrected *p* < .05; ** indicates corrected *p* < .01; *** indicates corrected *p* < .001.

Here are the temporal dynamics of WM representations for long-delay trials across the entire trial, including the encoding period. During the encoding period, no significant beta slope was found for either face or scene images. For statistics using Blocks 1 and 3, Face: *t*(19) = 1.39, *p* = .182, *d* = 0.31, *BF10* = 0.53. Scene: *t*(19) = -1.29, *p* = .212, *d* = 0.29, *BF10* = 0.48. For statistics using all three blocks, Face: *t*(19) = 1.10, *p* = .283, *d* = 0.25, *BF10* = 0.40. Scene: *t*(19) = -0.82, *p* = .424, *d* = 0.18, *BF10* = 0.31.

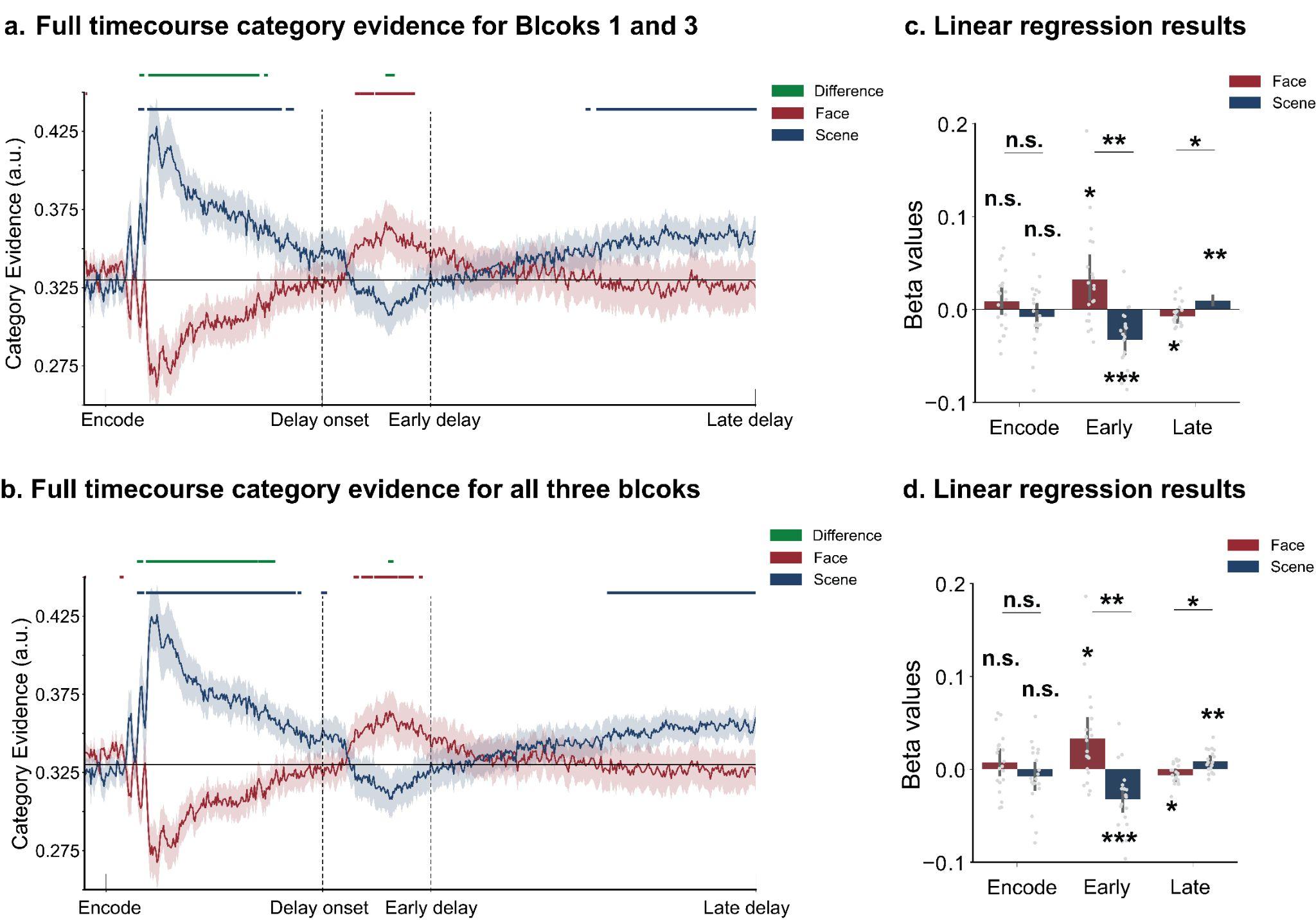

**Sfig 3.** **Temporal Fluctuations of Working Memory Representations across the full timecourse.** a). Category evidence for face and scene images for long-delay trials, excluding Block 2. b). Category evidence for face and scene images for long-delay trials, including Block 2. Red horizontal lines indicate time points where face evidence was significantly above the empirical chance level of 0.33; blue lines indicate significant scene evidence; green lines indicate time points where face and scene evidence differed significantly. The shaded area represents standard errors. c) & d). Beta values from linear regression models predicting category evidence from time, separately for the encoding, early (0–1 s), and late (1–4 s) delay windows. Error bars represent 95% confidence intervals. * indicates corrected *p* < .05; ** indicates corrected *p* < .01; *** indicates corrected *p* < .001; n.s. is not significant.

Here are the temporal dynamics of WM representations for short-delay trials across the entire trial, including the encoding period. For linear regression fits, we didn’t observe any significant results. For statistics using Blocks 1 and 3, Encoding period, Face: *t*(19) = 1.39, *p* = .182, *d* = 0.31, *BF10* = 0.53. Scene: *t*(19) = -1.29, *p* = .212, *d* = 0.29, *BF10* = 0.48. Difference between face and scene: *t*(19) = 1.43, *p* = .168, *d* = 0.60, *BF10* = 0.56. Early delay period: Face: *t*(19) = 0.60, *p* = .558, *d* = 0.13, *BF10* = 0.27. Scene: *t*(19) = -1.63, *p* =.120, *d* = 0.36, *BF10* = 0.72. Difference between face and scene: *t*(19) = -1.29, *p* = .212, *d* = 0.29, *BF10* = 0.48. For statistics using all three blocks, Encoding period, Face: *t*(19) = 1.10, *p* = .283, *d* = 0.25, *BF10* = 0.40. Scene: *t*(19) = -0.82, *p* = .424, *d* = 0.18, *BF10* = 0.31. Difference between face and scene: *t*(19) = 1.01, *p* = .323, *d* = 0.42, *BF10* = 0.37. Early delay period: Face: *t*(19) = 1.24, *p* = .232, *d* = 0.28, *BF10* = 0.45. Scene: *t*(19) = -2.28, *p* =.003, *d* = 0.51, *BF10* = 1.86. Difference between face and scene: *t*(19) = 1.86, *p* = .079, *d* = 0.77, *BF10* = 0.98.

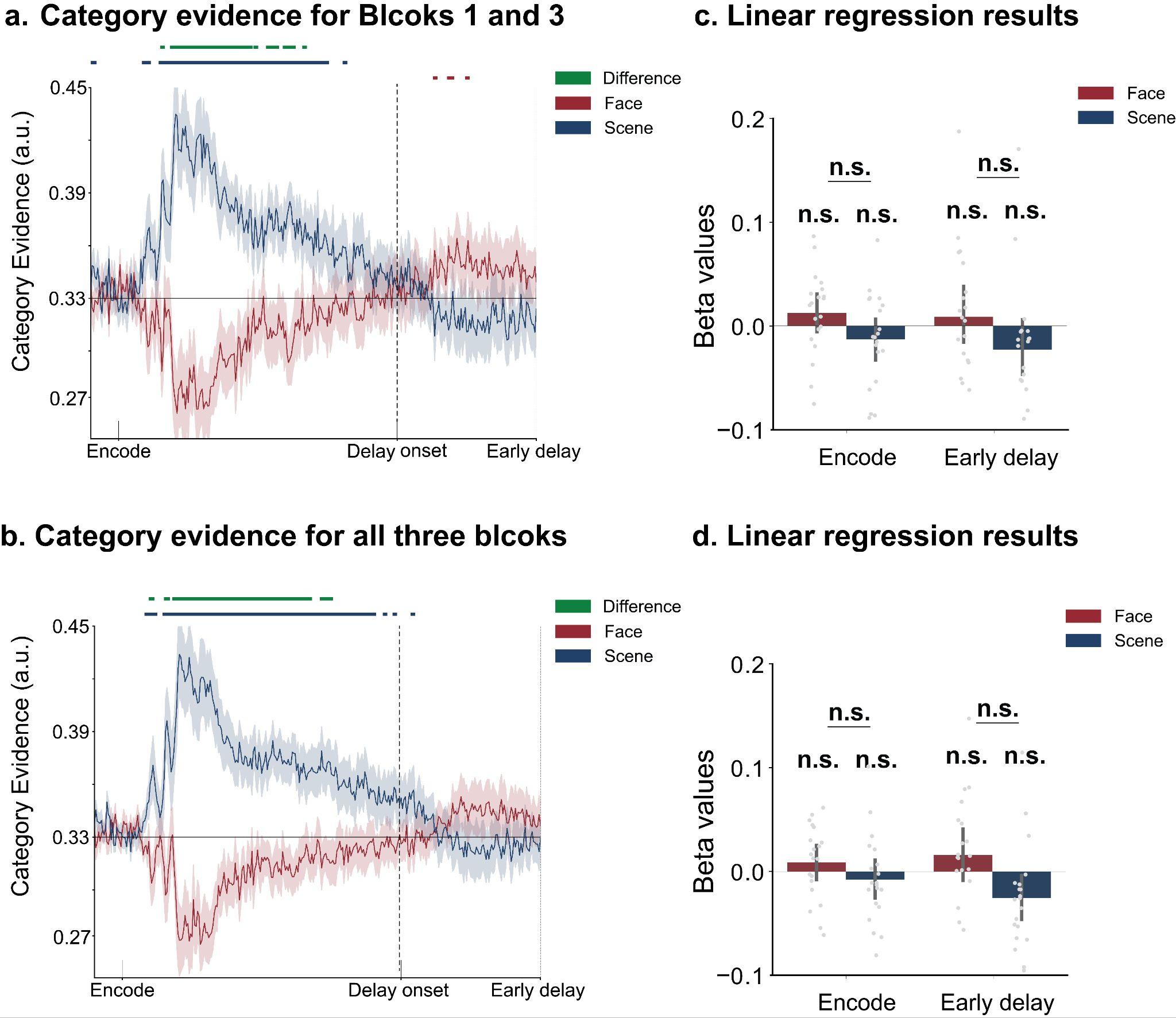

**Sfig 4.** **Temporal Fluctuations of Working Memory Representations for the short-delay trials.** a). Category evidence for face and scene images for short-delay trials, excluding Block 2. b). Category evidence for face and scene images for short-delay trials, including Block 2. Red horizontal lines indicate time points where face evidence was significantly above the empirical chance level of 0.33; blue lines indicate significant scene evidence; black lines indicate time points where face and scene evidence differed significantly. The shaded area represents standard errors. c) & d). Beta values from linear regression models predicting category evidence from time, separately for the encoding, early (0–1 s), and late (1–4 s) delay windows. Error bars represent 95% confidence intervals. n.s. is not significant.

Here are the results for linking LTM performance to representational fluctuations across the entire WM trial using Blocks 1 and 3, including the encoding period. During the encoding period, the category evidence difference for both remembered and forgotten trials was significantly above chance; however, no significant difference was observed between remembered and forgotten trials. Remembered: *t*(19) = 3.29, *p* = .004, *BF10* = 11.49. Forgotten: *t*(19) = 3.24, *p* = .004, *BF10* = 10.38. Remembered-Forgottern: *t*(19) = -1.45, *p* = .162, *BF10* = 0.58. Statistics for the early and late delay periods are shown in the main manuscript.

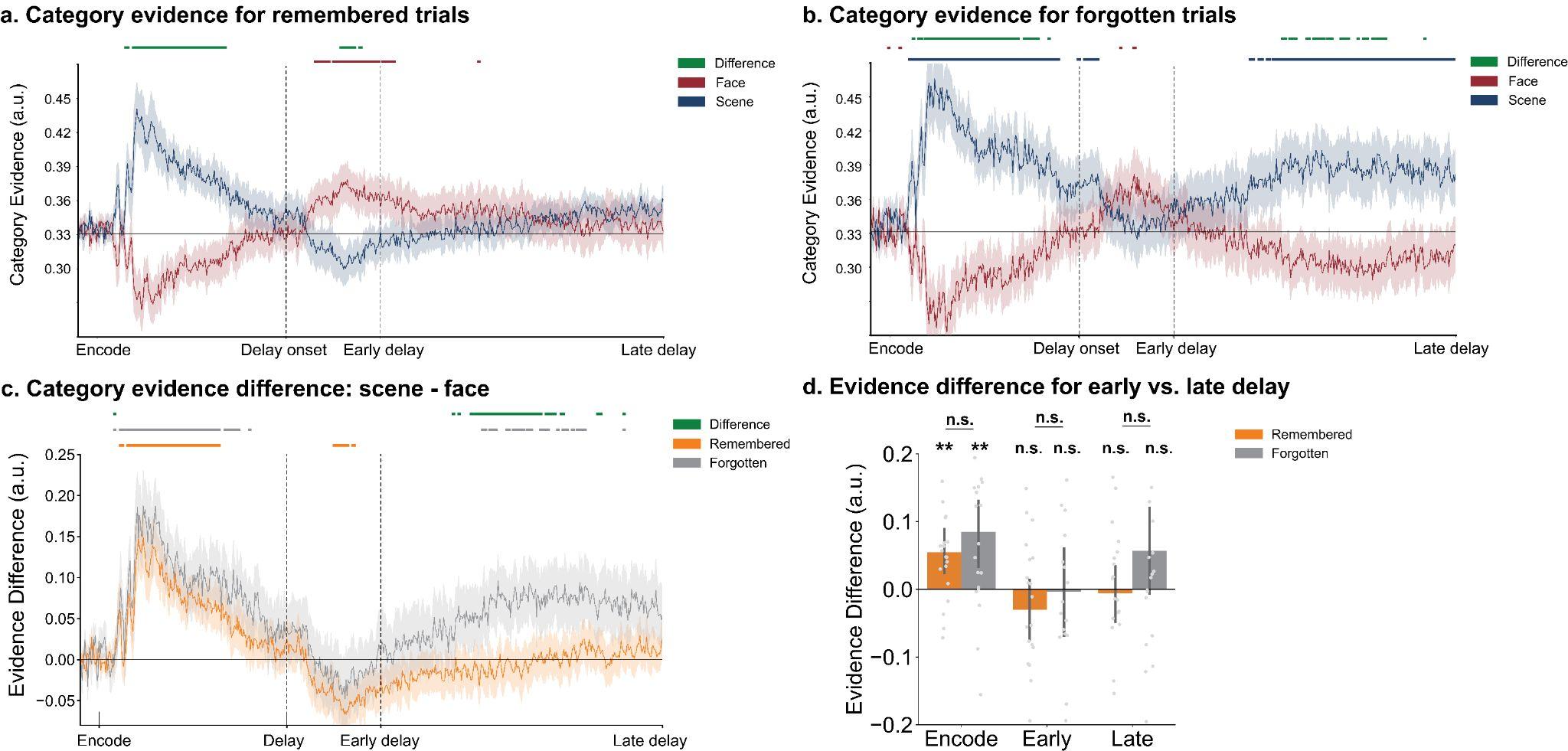

**Sfig 5. Differentiated temporal profiles for remembered and forgotten trials for Blocks 1 and 3 across the full timecourse.** a). Temporal dynamics of face and scene category evidence for trials in which face images were later remembered in LTM. b). Temporal category evidence for trials in which face images were later forgotten. Red horizontal lines indicate time points where face evidence was significantly above the empirical chance level (0.33); blue lines indicate significant scene evidence; green lines indicate significant differences between face and scene evidence. c). Differences in category evidence between face and scene for remembered and forgotten trials. Orange horizontal lines indicate time points where category evidence difference between face and scene was significantly above 0 for the remembered trials; grey lines indicate significant evidence difference for forgotten trials; green lines indicate significant differences between remembered and forgotten trials. The shaded area represents standard errors. d). Averaged category evidence difference for early (0–1 s) and late (1–4 s) delay windows. Error bars represent 95% confidence intervals. ** indicates corrected *p* < .01; n.s. is not significant.

Here are the results for linking LTM performance to representational fluctuations in WM using data from all three blocks. Similar to the results reported in the main text, we observed a larger category evidence difference between the face and scene in the forgotten trials than in the remembered trials during the late delay window (compare **Sfig. 6a** and **6b**). However, the effect was weaker, and fewer significant time points were identified when including data from Block 2 (**Sfig. 6c**). When averaging category evidence during the early and late delay periods, no significant difference was found between forgotten and remembered trials. Encoding period, Remembered: *t*(19) = 4.10, *p* < .001, *BF10* = 55.96. Forgotten: *t*(19) = 3.25, *p* = .004, *BF10* = 10.62. Remembered-Forgotten: *t*(19) = -0.97, *p* = .344, *BF10* = 0.35. Early delay period, Remembered: *t*(19) = -1.09, *p* = .287, *BF10* = 0.39. Forgotten: *t*(19) = -0.67, *p* = .511, *BF10* = 0.28. Remembered-Forgotten: *t*(19) = -0.97, *p* = .342, *BF10* = 0.35; Late delay period, Remembered: *t*(19) = 0.04, *p* = .969, *BF10* = 0.23. Forgotten: *t*(19) = 1.10, *p* = .284, *BF10* = 0.40. Remembered-Forgotten: *t*(19) = -2.15, *p* = .045, *d* = 0.89, *BF10* = 1.52.

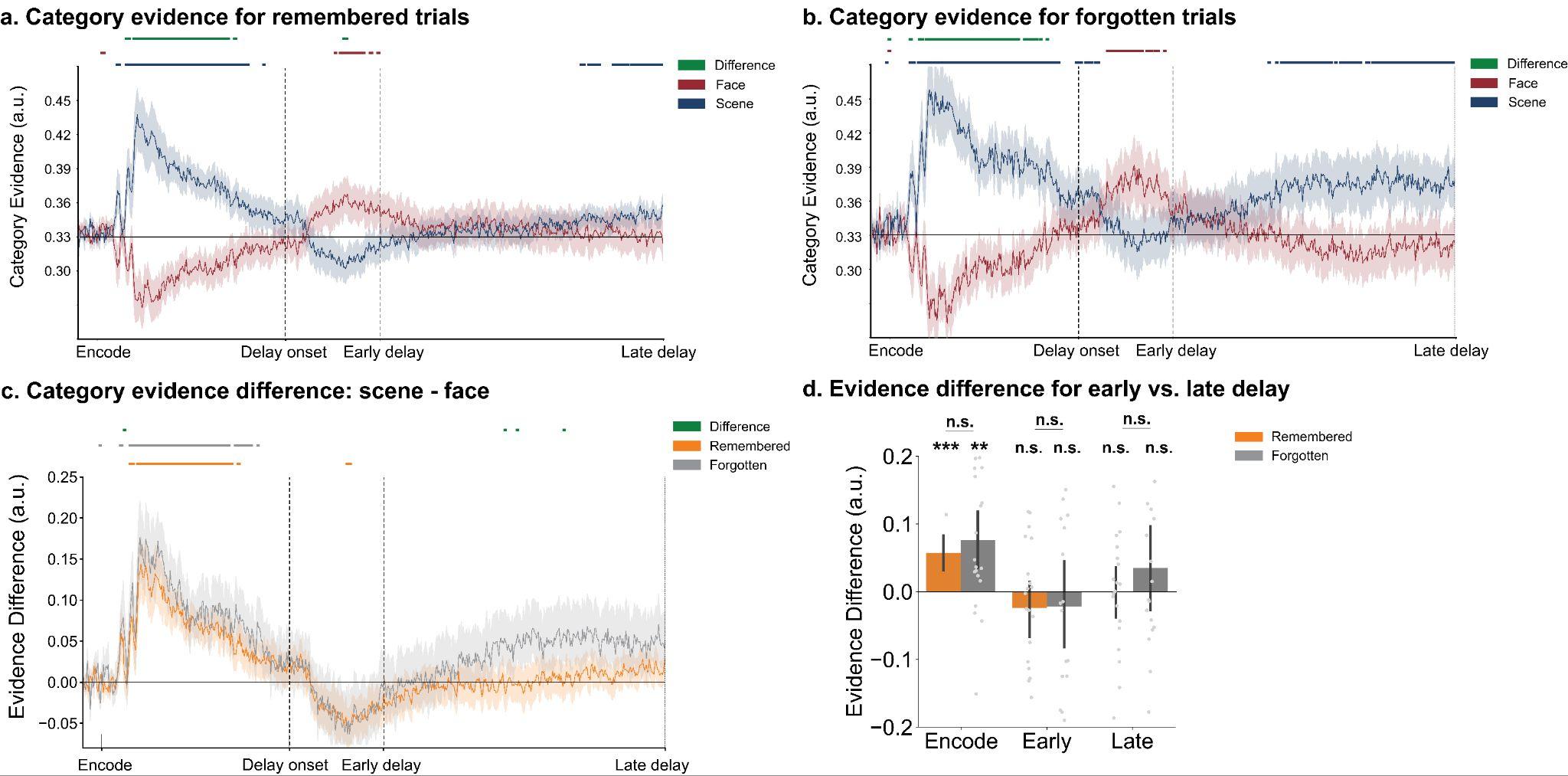
**Sfig 6.** **Differentiated temporal profiles for remembered and forgotten trials for all three blocks across the full timecourse.** a). Temporal dynamics of face and scene category evidence for trials in which face images were later remembered in LTM. b). Temporal category evidence for trials in which face images were later forgotten. Red horizontal lines indicate time points where face evidence was significantly above the empirical chance level (0.33); blue lines indicate significant scene evidence; green lines indicate significant differences between face and scene evidence. c). Differences in category evidence between face and scene for remembered and forgotten trials. Orange horizontal lines indicate time points where category evidence difference between face and scene was significantly above 0 for the remembered trials; grey lines indicate significant evidence difference for forgotten trials; green lines indicate significant differences between remembered and forgotten trials. The shaded area represents standard errors. d). Averaged category evidence difference for early (0–1 s) and late (1–4 s) delay windows. Error bars represent 95% confidence intervals. ** indicates corrected *p* < .01; *** indicates corrected *p* < .001; n.s. is not significant.

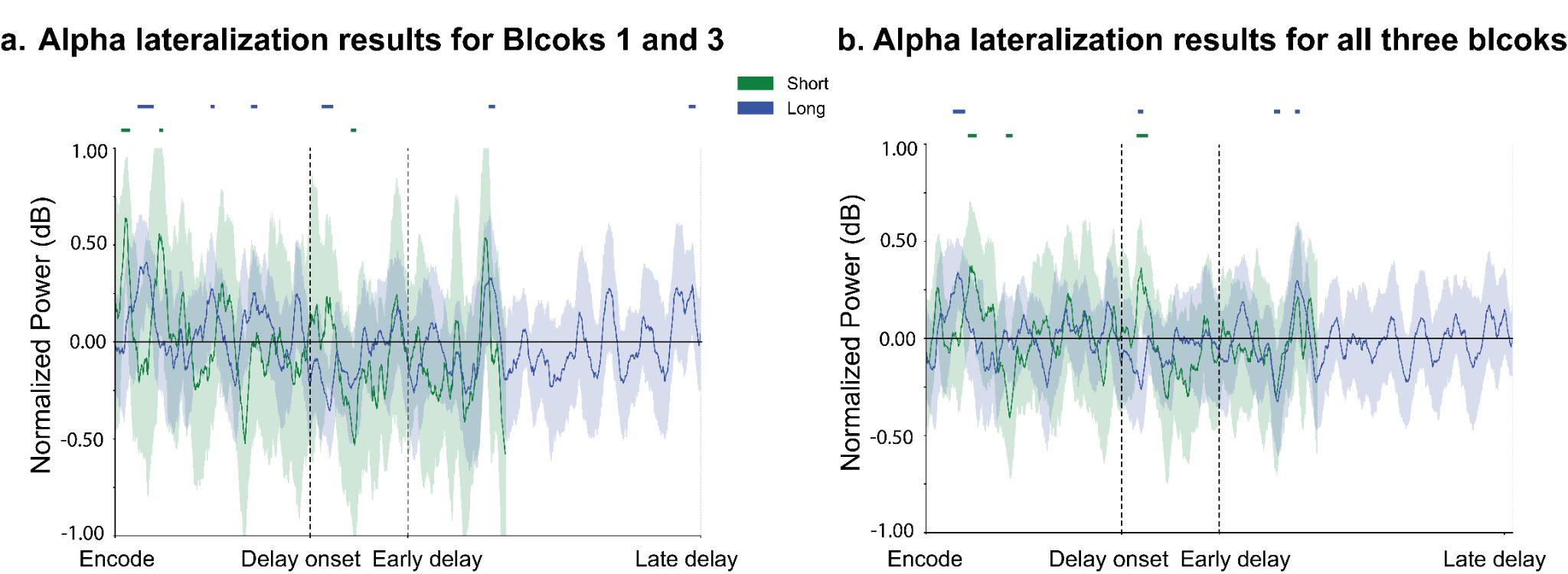

**Sfig 7. Lateralized alpha results.** a). Lateralized alpha power using data from Blocks 1 and 3, excluding Block 2. b). Lateralized alpha power using data from all three blocks, including Block 2. Green horizontal lines indicate time points where lateralized alpha power is significantly different than 0 for short-delay trials, while blue horizontal lines indicate significant time points for long-delay trials. The shaded area represents standard errors.
